# A bird’s-eye view: Evaluating drone imagery for the detection and monitoring of endangered and invasive day gecko species

**DOI:** 10.1101/2023.03.20.533518

**Authors:** Nicolas Dubos, Xavier Porcel, Markus A. Roesch, Juan Claudin, Romain Pinel, Jean-Michel Probst, Gregory Deso

## Abstract

Herpetofauna monitoring can be strongly limited by terrain accessibility, impeding our understanding of species ecology and thus challenging their conservation. This is particularly true for species living in the canopy, on cliffs or in dense vegetation. Remote sensing imagery may fill this gap by offering a cost-effective monitoring approach allowing to improve species detection in inaccessible areas. We investigated the applicability of drone-based monitoring for a Critically Endangered insular gecko (*Phelsuma inexpectata*) and two invasive alien species representing a risk for the former (*P. grandis* and *P. laticauda*). We determined the approach distance before inducing behavioural response caused by the drone’s presence. All three study species showed no reaction to the drone’s presence until very close distances (mean distance for *P. inexpectata*: 33.8 cm; *P. grandis*: 21.9 cm; *P. laticauda*: 26.4 cm). We then performed horizontal and vertical approaches, taking photos every meter starting at 10 m away from the canopy edge to determine an optimal distance for detection while ensuring species-level identification. We examined a total of 328 photos. We found a bimodality in the number of detected geckos, with different individuals recorded between short and intermediate distances. Therefore, we recommend taking photos at two distances of 2–2.5 m and 5 m away from the canopy, ideally facing away from the sun and in low wind conditions. We encourage the application of our methodology for *Phelsuma* spp., but also for other species of similar size and ecology to improve detection in inaccessible areas.

## Introduction

Population assessment of endangered and rare species are often limited by a multitude of factors, including funding, observer experience, species detectability and terrain accessibility. Consequently, species living in complex habitats or with dynamic habitat use are poorly understood and their conservation becomes challenging. The development of novel cost-effective methods for cryptic or threatened species monitoring is a priority for their conservation (Monks, Wills, and Knox, 2022). Insular endemic species are particularly vulnerable to invasive alien species (IAS). The spread of IAS is generally facilitated by lack of surveillance efforts, preventing their early detection and allowing for initial dispersal (Cuthbert et al., 2022). The rising economic costs of IAS encourage the settlement of preventive measures (Diagne et al., 2021). Detection and monitoring of IAS may be considerably improved with the development of cost-effective monitoring methods.

Recent technological advancements and reduced costs for electronic devices have contributed to the development of novel methods for biodiversity monitoring. Novel methodologies such as computer assisted slow-speed road cruising (Jones et al., 2022) or camera-trapping (e.g., Roesch, Hansen, and Cole, 2021; Deso, Crouzet, and Bonnet, 2022) have already proven efficient. Drones have recently been used for the monitoring of endangered species (Landeo-Yauri et al., 2020; Varela-Jaramillo et al., 2023), including cryptic reptiles (Monks, Wills, and Knox, 2022), and have improved the detection of invasive reptile species living in the tree canopy (Aota et al., 2021).

The Critically Endangered Manapany day gecko, *Phelsuma inexpectata* Mertens 1966, is endemic to Reunion Island. Its distribution is restricted to a narrow stripe along the southern coastline. The species frequently uses screw pine *Pandanus utilis,* where it can be locally abundant (Bour, Probst, and Ribes, 1995). Its habitat use is dynamic throughout the seasons, with a more frequent use of the canopy during winter (Choeur et al., 2023). The development of a year-round remote sensing monitoring protocol dedicated to this species may increase detection, improve the temporal resolution of surveys, helps understanding the species’ ecology and ultimately improves conservation management.

Among the few fragmented populations of *P. inexpectata*, several have been reported in sympatry with invasive *Phelsuma* spp., i.e. the Madagascar giant day gecko *P. grandis* Gray 1870 and the gold dust day gecko *P. laticauda* Boettger 1880 (Dubos, 2013; Porcel et al., 2021). A colonisation of Manapany-les-Bains, the stronghold of *P. inexpectata*, by *P. grandis* has successfully been controlled by the NGO Nature Océan Indien between 2010–2012 and *P. grandis* has not been observed in the area ever since (M.A. Roesch pers. obs.). Both invasive species can thrive in similar habitats to *P. inexpectata* and share resources, inducing competition (Hoarau et al., 2021; Deso et al., 2023; Porcel, Luspot, and Probst, 2023). *Phelsuma grandis* also raises concerns due to its larger size, imposing high predation risk on smaller species (Buckland et al., 2014). Both invasive species successfully established throughout the world (Dubos et al., 2014; Fieldsend and Krysko, 2019; Fieldsend, Borgia, and Krysko, 2020; Fieldsend et al., 2021; Dubos et al., 2022a), with strong invasion potential on tropical islands (Dubos et al., 2022a). The two invasive *Phelsuma* spp. can be found in a variety of habitats including primary forests, shrub land, urban environment and agricultural areas (D’Cruze et al., 2009; Dubos et al., 2014). Beyond promoting early detection in uninvaded areas, the use of remote sensing may help understanding their impact on native species where they are already established. Drone imagery offers a bird’s-eye view on areas that are otherwise inaccessible or difficult to survey. It can improve species detection and thus, contribute to the monitoring and spread of IAS. It may also allow for the study of interactions between native species and IAS and to better characterize the dynamics of habitat use in areas invisible to the observer on the ground.

This study investigates the use of remote sensing-based monitoring of native and invasive *Phelsuma* spp., with the aim to improve detection probability in an otherwise inaccessible area: tree canopy. We (i) quantified the behavioural response of geckos to the approaching drone, (ii) determined the optimal distance for maximum detection and (iii) investigated variation in detection relative to time of day and species-level identification. We eventually propose a standardized framework for the monitoring of *Phelsuma* spp. based on drone imagery.

## Methods

### Study sites

Our research took place at three sites on Reunion Island: (1) in the village of Manapany-les-Bains (−21.37 S; 55.58 E; conducted on 22/11/2022), at a site where only *P. inexpectata* is present; (2) in the botanical garden *Domaine du Café Grillé* (−21.37 S; 55.42 E; conducted on 23/11/2022) where *P. inexpectata* and *P. laticauda* co-occur; (3) in a public park in the city of Saint Benoît (−21.03 S; 55.72 E; conducted on 25/11/2022) occupied by *P. grandis.* In all three sites, surveys were conducted along screw pines, *Pandanus utilis,* which represent a highly favourable habitat for either species.

### Material

We used a DJI Phantom 4 Pro V2.0 drone equipped with its standard camera. The camera has a 1-inch 20M pixel sensor and a 24 mm (35 mm format equivalent) lens, corresponding to an 84° field of view. All take-offs and landings were located in secured and open areas, with restricted access to the public, and at least 10 m away from the geckos’ habitat.

### Determining approach distance

We tested whether the presence of a drone would induce a behavioural response in our three study species. We first located individuals which could be approached safely by the drone until a short distance based two criteria: (1) no obstacle between the drone and the gecko and (2) little canopy cover for precise drone geolocation and manoeuvrability.

We stabilised the drone image at 10 m distance from the monitored individual at its height. Then, we steadily flew the drone horizontally towards the individual. We interrupted the approach either when the individual reacted to the drone’s presence (i.e. when observing an escape behaviour), or when the individual was about 20 cm away from the drone propellers (for the individual’s safety and material integrity). Therefore, a distance of 20 cm suggests that the individual did not respond to the drone’s presence.

The drone may induce a different impact on the target species depending on the approach orientation (e.g., perception of avian predator and potential effect of propellers’ blow). We used the aforementioned method to evaluate the vertical approach distance for *P. inexpectata, s*ince this species is frequently observed on the ground (mostly on volcanic rock beaches; Deso and Probst, 2007).

#### Statistical analysis

Since insular species are known for having lost vigilance regarding predators, we expected the two invasive species to respond to the drone at longer distances than *P. inexpectata*. We tested whether the approach distance would differ between species with a linear model (LM, assuming a gaussian distribution). We removed the data related to vertical approaches, since such data could only be acquired for *P. inexpectata*. We used the distance of approach as the response variable and the species as explanatory variable.

For *P. inexpectata*, we expected a stronger response in the vertical approach because they are known to respond to bird predators, such as the Reunion harrier *Circus maillardi* and the red-whiskered bulbul *Pycnonotus jocosus* (J.-M. Probst pers. obs.). We built a second LM with approach distance as the response variable and approach orientation as predictor. We expected differences in the response to the drone between adult and juvenile geckos, thus added to the model the maturity of individuals as a two-level factor effect (Adult *versus* Juvenile).

### Determining optimal detection distance

We performed horizontal and vertical approaches. For horizontal approaches, we stabilized the drone at the canopy level, i.e. between three and six meters above ground (depending on tree height) and at a horizontal distance of 10 m from the canopy, with the camera oriented in opposite direction to the sun when applicable. We flew the drone steadily towards the tree and took photos every meter until reaching a distance of 1 m.

For vertical approaches, we first measured the canopy height with the drone embedded barometer and GPS, then started approaching from 10 m above the canopy. We repeated the operation four times between 8:00am and 2:00pm. At the shortest distances, where the camera’s field of view could not enable us to encompass the whole tree, we took multiple photos at the same distance to cover the entirety of the canopy. Images were carefully examined by three observers afterwards (GD, ND, XP), with three to five minutes of effort per photo depending on image complexity. During each drone operation, we performed a standardised point count survey (human visual counts) with two to three observers (JC, ND, XP) per site. We counted all visible geckos up to a distance of 8 m from the observer with an increment of 2 m, resulting in four increments per count for a duration of one minute per increment.

#### Statistical analysis

We used a Generalized Additive Mixed Model (GAMMs; R package mgcv version 1.8-42; Wood, 2011) assuming a Poisson distribution with gecko count as the response variable and drone distance as spline effect to examine variation in gecko detection on images. For both drone image and human visual counts, we first performed the analysis for all species combined. Models included a species and an observer categorical fixed effect, and a sampling session random effect. We then repeated the analysis for each species individually, accounting for the effect of observer (fixed effect) and a sampling session (random effect). We added a site effect for *P. inexpectata*, because this species was observed at two sites.

### Assessing time of day effect on detection and distance on species-level identification

We examined whether there was an optimal time of day to maximize detection during the four drone sessions performed between 8:00am and 2:00pm described above. We used a Generalized Additive Model (GAM; Poisson family), with gecko count as the response variable and time of day as spline effect. We accounted for differences in species abundance with a species adjustment variable.

Eventually, we assessed the maximum distance for a species-level identification using a GAMM (Poisson family) to predict the effect of distance (spline effect) on unidentified species count (response variable). We added an observer effect as a fixed effect and sampling session as a random effect. All analyses were performed under R version 4.1.3 (R Core Team, 2022).

## Results

### Determining approach distance

We measured the approach distance for 26 individuals (*P. inexpectata n* = 11; *P. grandis n* = 8; *P. laticauda n* = 7), including 19 adults and 7 juveniles. Interestingly, we found overall very little effect of the drone’s presence on all three study species (fig. 1). The approach distance to *P. inexpectata* was significantly different from zero (mean ± SE = 33.8 cm ± 5.4; *P* = 0.02), while it did not significantly differ for the two IAS (*P. grandis* mean ± SE = 21.9 cm ± 4.7; *P* = 0.70; *P. laticauda* mean distance ± SE = 26.4 cm ± 5.0; *P* = 0.22).

**Figure 1.**
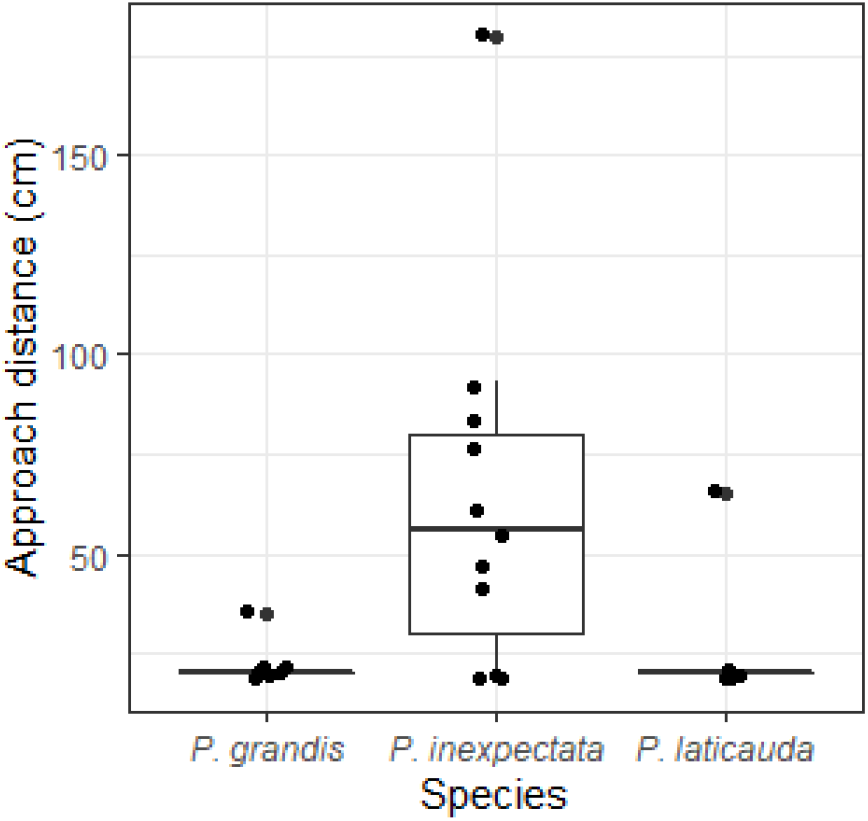
Approach distance before the drone induced behavioural response for three *Phelsuma* species at two stages of maturity (total = 26). Boxes represent the first and third quartiles, the horizontal bar represents the median and the points represent outliers. We show jittered data points.

We found a significant difference between horizontal and vertical approach distances for *P. inexpectata* (table 1). As expected, the approach distance was longer when approaching vertically (+37.3 cm). We found no statistical effect of maturity.

**Table 1.** Model estimates for the effect of approach orientation and maturity on the approach distance before behavioural response to the drone’s presence for *Phelsuma inexpectata*. The significant effect is shown in bold.

|  | Estimate | SE | *P* |
| --- | --- | --- | --- |
| Intercept (Adult, Horizontal) | 20.75 | 17.24 | 0.26 |
| Maturity (Juvenile) | -20.75 | 29.87 | 0.51 |
| **Orientation (Vertical)** | **58.05** | **23.14** | **0.03** |

### Determining optimal detection distance

We produced and examined a total of 328 drone photos. We counted between 0 and 6 *P. inexpectata* per sampling unit (mean ± SD = 0.70 ± 1.20 at given distance, sampling session, site and observer group; fig. 2) on the drone images. With human visual counts we counted between 0 and 9 individuals per sampling unit (mean ± SD = 2.33 ± 2.37). For *P. laticauda,* we counted between 0 and 6 (mean ± SD = 0.57 ± 1.00), and between 4 and 19 (mean ± SD = 10.40 ± 4.49) individuals per sampling units, respectively for both methods. For *P. grandis*, we counted between 0 and 1 (mean ± SD = 0.09 ± 0.29) and between 0 and 3 (mean ± SD = 0.93 ± 1.03) individuals per sampling units, respectively.

**Figure 2.**
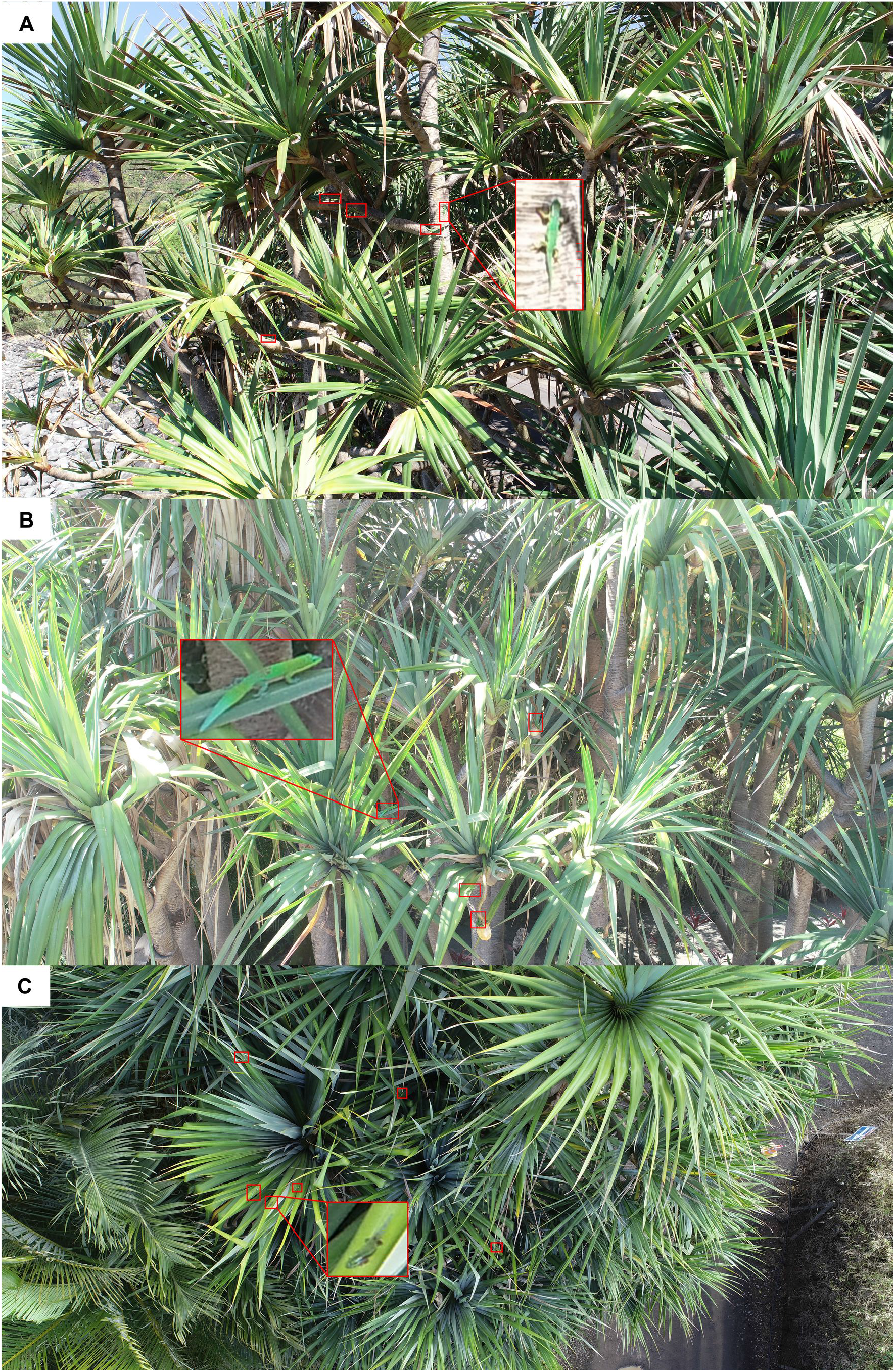
Drone images of *Phelsuma inexpectata* (A) and *P. laticauda* (B) on horizontal approach, and *P. inexpectata* (C) on vertical approach. Individuals are highlighted with red rectangles.

The field of view strongly differed between short and long distances (e.g. 2 m *versus* 5 m). We found that different individuals may be detected within the same sampling session depending on the angle and field of view. We assume individuals were different based on their different location between short time intervals, and difference in size or sex.

We identified two modalities in gecko detection with drones with all species combined (fig. 3). The highest detection rates were at 2.5 m and 5.5 m distance. The detection of *P. inexpectata* increased until reaching a first plateau near 5 m, then further increased between 4 and 2 m before reaching a second plateau (fig. 4). The highest detection rate was between 2 and 6 m for *P. laticauda.* Detection decreased linearly with the distance for *P. grandis*. Detection with the human visual counts approach decreased linearly with distance in all three species (fig. 4).

**Figure 3.**
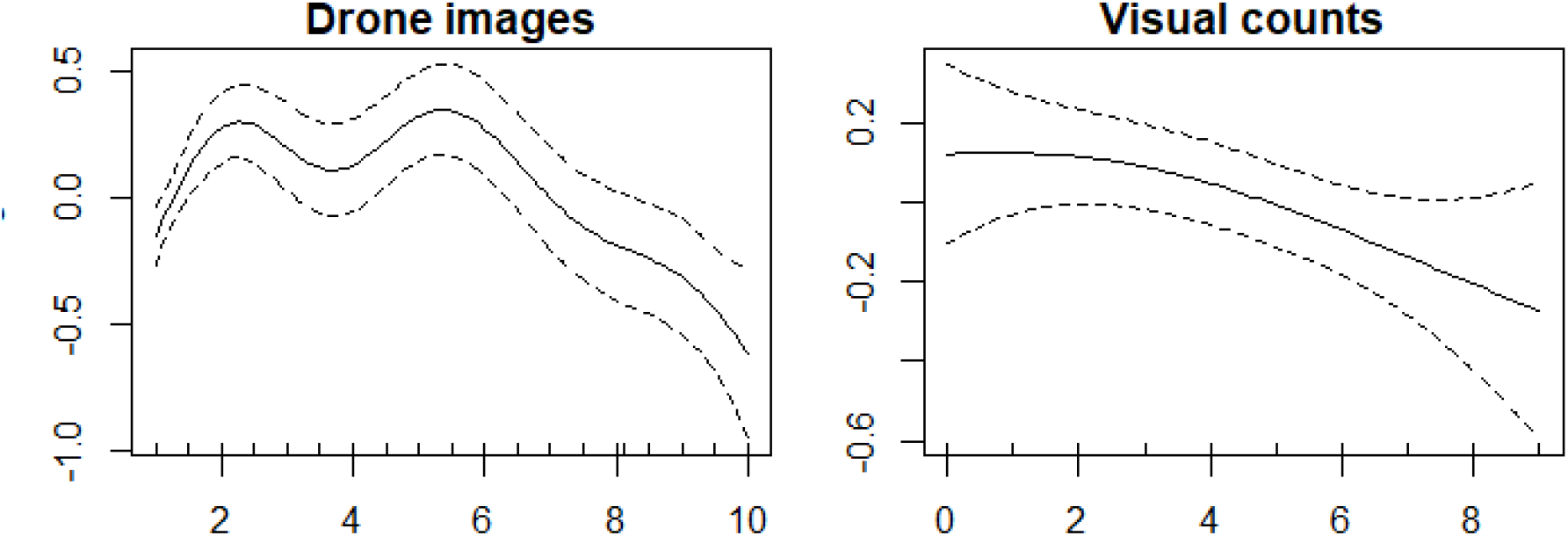
Effect of distance on gecko detection (predicted values obtained from GAMMs, three *Phelsuma* species combined) with two methods of observation (left: drone imagery; right: human visual counts).

**Figure 4.**
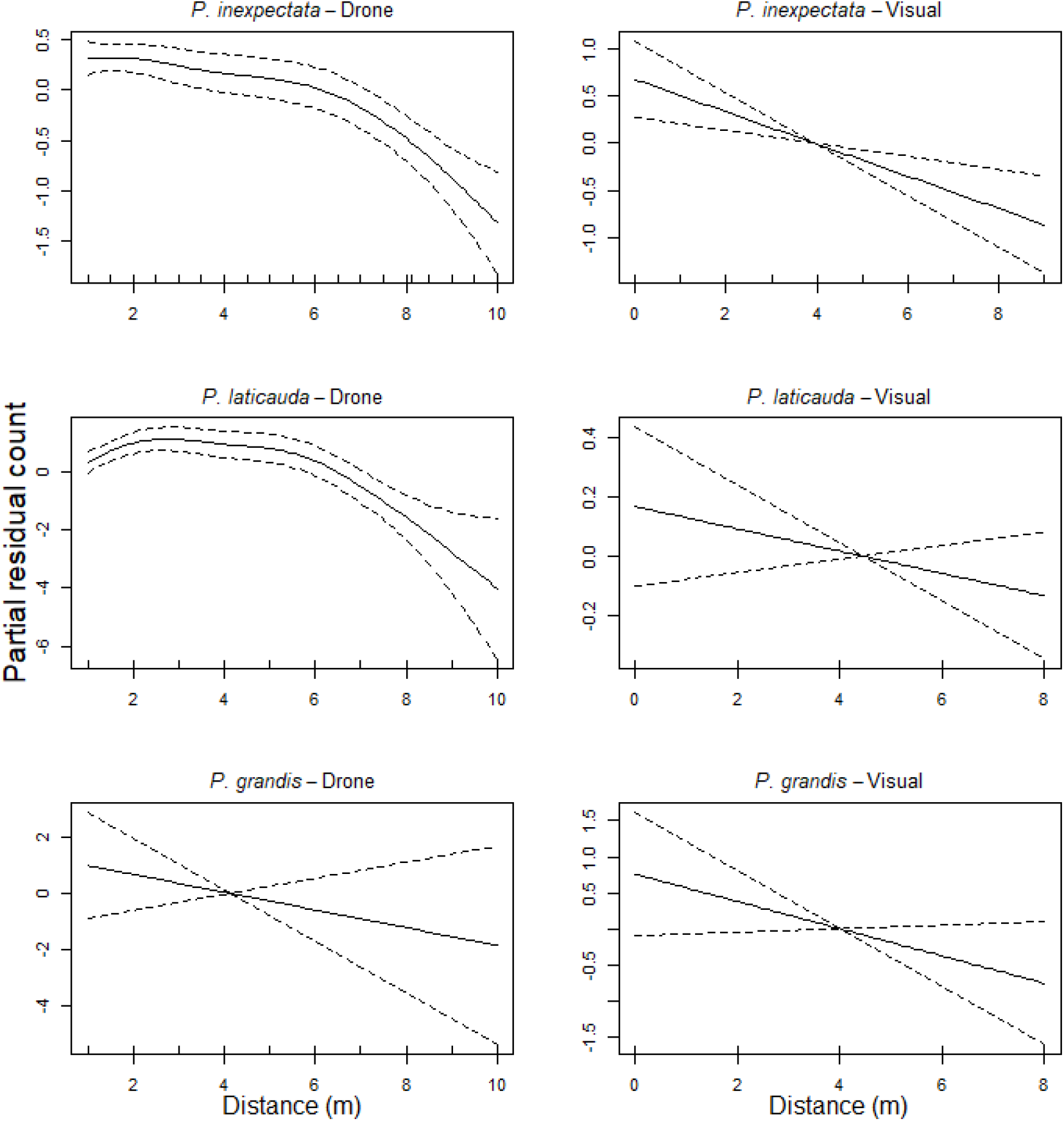
Effect of distance on gecko detection for three *Phelsuma* species (predicted values obtained from GAMMs) with two methods of observation (left: drone imagery; right: human visual counts).

### Determine time of day effect on detection and distance on species-level identification

The number of geckos detected was stable throughout the morning but became more variable at around 11:00am, and eventually decreased linearly after 12:00pm (fig. 5). Species-level identification was low at a distance between 10 m and 6 m, then the rate of unidentified species decreased as the drone approached (fig. 5).

**Figure 5.**
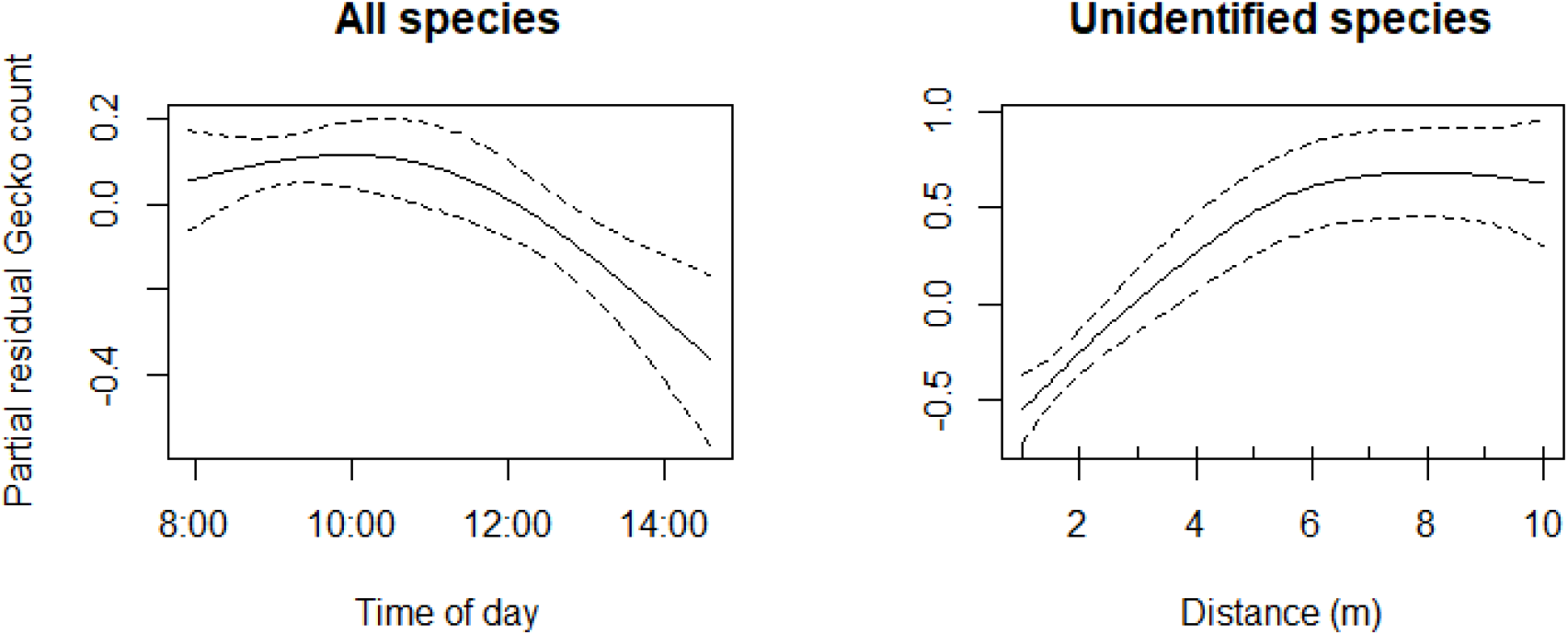
Variation in the number of geckos detected depending on time of day (left panel) and variation in the number of unidentified species through distance (right panel).

## Discussion

Drone imagery is a promising avenue for the monitoring of *Phelsuma* species. Our three study species showed very little behavioural response to the drone’s presence, and drone images enabled us to detect many individuals in the canopy, which otherwise remained undetected by eye (e.g., fig. 2 C). Approach distances were unexpectedly short, even shorter than previously found in New Zealander lizards, with 33.8 cm in average for *P. inexpectata versus* approximately 59 cm for the Jewelled geckos *Naultinus gemmeus* and 107 cm for the grand skinks *Oligosoma grande* (Monks, Wills, and Knox, 2022). This allowed for short-distance photo taking and high-resolution imagery.

In accordance with Varela-Jamarillo et al. (2023), geckos were less disturbed by the drone than by human presence at the same distance, suggesting that our approach is non-invasive. We showed that *P. inexpectata* was more sensitive to vertical approaching. This is possibly due to the conservation of anti-avian predator behaviour for bird species which were among the few native predators before human settlement in Reunion Island. Overall, all three species showed little behavioural response and allowed close drone encounters. The native *P. inexpectata* reacted more than the two exotic ones. This might be unexpected because oceanic island species have lost anti-predator behaviours (Blumstein et al., 2005), and Madagascar is considered a continental island (Andreone et al., 2021). This suggest that our method can be applied to other *Phelsuma* species (e.g. the more cryptic *P. borbonica* in Reunion Island), both in oceanic islands and continental systems.

Human visual counts resulted in the detection of more individuals than using the drone. This is presumably due to our choice of study site, with very accessible trees with good visibility on the tree trunks (which constitute important supports for thermoregulation in *Phelsuma* geckos). However, the use of drone imagery was highly complementary to the visual counts, since we detected additional individuals on *Pandanus* leaves in the canopy. We assume that the benefit of drone-based monitoring might become clearer in less accessible areas such as cliffs and shrublands and may outperform visual counts (Monks, Wills, and Knox, 2022; Varela-Jaramillo et al., 2023). *Phelsuma inexpectata* is distributed along the coastline, inhabiting steep slopes and cliffs. A comprehensive survey performed throughout the whole distribution of *P. inexpectata* showed important spatial gaps in sample sites due to accessibility (Dubos, 2010), which could be filled with our approach. Future sampling effort may be oriented towards these remnant natural habitats and other unprospected areas to identify potential new populations. Similarly for the two invasive species, which are more likely to disperse through the dense vegetation, drone-based surveys may improve the current knowledge of their distribution and help monitor their spread (Aota et al., 2021). At one of our study sites (the botanical garden *Domaine du Café Grillé), P. laticauda* and *P. inexpectata* co-occur. This area and its surroundings were predicted as hosting the most suitable climate in the future for the endemic *P. inexpectata* (Dubos et al., 2022b). On the other hand, climate change is predicted to benefit *P. laticauda* (Dubos et al., 2022a), which emphasizes the need to pursue the sampling effort at this site in order to better understand the impact of the invasive *P. laticauda* on the Critically Endangered *P. inexpectata* and plan efficient intervention if needed.

### Methodological recommendations

Drone-based monitoring should be carried out at the height corresponding to the upper part of the canopy when wind conditions are favourable. When applicable, the camera should orientated in opposite direction to the sun to avoid backlight and because geckos are frequently observed on sun spots for thermoregulation. We found a bimodality in detection rates with all species combined (but not in species-specific models, presumably because larger sample size allowed higher degrees of freedom for the spline effect), with different individuals identified between modalities. Therefore, we recommend taking two photos respectively at a distance of 2–2.5 m and 5 m, both horizontally and vertically. For large trees at short distances (2–2.5 m), multiple photos may be taken in order to cover the whole canopy. Photos at 5 m distance offer a fair trade-off between field of view (encompassing more vegetation) and image resolution for species-level identification. Photos taken at 2 m were highly complementary since they benefit from a higher resolution and a sufficiently different angle to allow the detection of different individuals and more accurate species identification. Photos taken at shorter distances may provide too narrow field of view, hence the fewer geckos detected in the present study. These distance recommendations stand for a medium size drone and a camera with similar specifications to those used in this study (1-inch 20M pixel sensor and 24 mm lens), and may be adjusted should the drone and camera differ much from these characteristics.

### Concluding remarks

Remote sensing-based survey offers the opportunity to improve detection in inaccessible areas, increases the temporal resolution of *Phelsuma* spp. monitoring and eventually develop automated artificial intelligence-based gecko detection. The use of deep learning techniques has already proven efficient in the monitoring of invasive arboreal lizards of similar size to our *Phelsuma* spp. (i.e. *Anolis carolinensis;* Aota et al., 2021) and may be also developed for our context. This offers the opportunity to develop proactive surveillance programmes, hence improve the chances of early detection and eventually help in the reduction of the impact of invasive species.

We showed that species-level identification was reliable within 5 m distance from the geckos. However, this approach may not be suitable for individual-level identification with the current resolution of standard mid-range drone cameras and may only be possible for larger species (e.g., photo-identification of Galàpagos marine iguanas; Varela-Jaramillo et al., 2023). Further improvement of mid-range drone camera lenses in the future might allow for higher resolution imagery and thus, individual identification.

The habitat use of *P. inexpectata* is dynamic, with more frequent use of the canopy during winter (Choeur et al., 2023). Our survey was carried out in summer, and we therefore expect better detection rates during winter. Future surveys should be performed throughout the year for a better understanding of habitat use dynamics of the species. This aspect also needs to be explored for the two invasive species using the same methodology. This will enable researchers and operators to increase the spatial coverage and the cost-effectiveness of surveillance efforts. We encourage the application of our methodology for *Phelsuma* spp. monitoring and other species, either endangered or invasive ones, of similar size and ecology throughout the world.

## Acknowledgements

We would like to thank all the staff of the botanical garden *Domaine du Café Grillé* for their support. We thank Annelise Tran for her suggestions and Lucas Grosolia for his assistance in the field. All flights were performed abiding by the French drone laws and regulations and all permissions were obtained prior surveys in private properties.

